# Light-induced asymmetries in the embryonic retina are mediated by the vascular system and extracellular matrix

**DOI:** 10.1101/2021.10.08.463575

**Authors:** Elisabetta Versace, Paola Sgadò, Julia George, Jasmine L. Loveland, Joseph Ward, Peter Thorpe, Lars Juhl Jensen, Karen A. Spencer, Silvia Paracchini, Giorgio Vallortigara

## Abstract

Left-right asymmetries in the nervous system (lateralisation) influence a broad range of behaviours, from social responses to navigation and language. The role and pathways of endogenous and environmental mechanisms in the ontogeny of lateralisation remains to be established. The domestic chick is a model of both endogenous and experience-induced lateralisation driven by light exposure. Following the endogenous rightward rotation of the embryo, the asymmetrical position in the egg results in a greater exposure of the right eye to environmental light. To identify the genetic pathways activated by asymmetric light stimulation, and their time course, we exposed embryos to different light regimes: darkness, 6 hours of light and 24 hours of light. We used RNA-seq to compare gene expression in the right and left retinas and telencephalon. As expected, no differential expression between left and right was present in darkness. We detected differential gene expression in right vs left retina after 6 hours of light exposure. This difference disappeared before 24 hours of light exposure, suggesting that light-induced activation is a self-terminating phenomenon. This transient effect of light exposure was associated with a downregulation of the sensitive-period mediator gene DIO2 (iodothyronine deiodinase 2) in the right retina. No differences between genes expressed in the right vs. left telencephalon were detected. Gene networks associated with lateralisation were connected to vascularisation, cell motility, and the extracellular matrix. Interestingly, we know that the extracellular matrix– including the differentially expressed *PDGFRB* (platelet-derived growth factor receptor β) gene – is involved in both sensitive periods and in the endogenous chiral mechanism of primary cilia, that drives lateralisation. Our data show a similarity between endogenous and experience-driven lateralisation, identifying functional gene networks that affect lateralisation in a specific time window.

## Introduction

Left-right asymmetries (lateralisation) are a major principle of organization of the nervous system in both vertebrate and invertebrate species [1–4]. Anatomical and functional lateralisation has a central role in behaviour and cognition from foraging to navigation, limb use, attention, communication, social responses, and language [1,3–6]. Lateralisation often provides increased neural and cognitive efficiency [1,3,7–12], whereas atypical lateralisation is associated with disease [13,14]. Despite the importance of lateralisation, its ontogenesis is far from being understood. Endogenous, genetically-guided mechanisms drive the emergence of internal organ asymmetries [15]. Endogenous mechanisms, though, do not explain the whole ontogenesis of lateralisation [11,16,17], or the dissociation of lateralisation patterns observed across different tissues [6,18,19]. Here we focus on the ontogeny of experience-driven lateralisation and its connection to endogenous lateralisation, using domestic chicks as a model system.

Endogenous mechanisms have been identified as factors that initiate systematic left-right asymmetries [20,21]. In particular, mounting evidence shows that left-right asymmetry in embryonic development is mediated by cilia [22]. Cilia are hair-like cell organelles that extend from the cell surface into the extracellular space [14,23,24]. It has been shown that cilia play an important role in establishing the left-right body axis. Due to molecular chirality, motor cilia induce a leftward flow of extracellular fluid detected by primary (or sensory) cilia [25]. These activate stronger left-sided Ca^2+^ signalling [26,27], stronger left-sided expression of the Nodal pathway [13,reviewed in 14], and in turn left-right asymmetries. Defects of cilia include the lateralised *situs inversus* phenotype, where the major visceral organs appear flipped compared to the typical position [28]. While the role of cilia in the asymmetries of the central nervous system is not entire clear, more than a hundred human disorders have been linked to defects in cilia [29].

Beside endogenous mechanisms, environmental/experience-related factors [30,31] and epigenetic factors [31–33] affect lateralisation at the anatomical and functional levels. The role of experience-related factors in the ontogenesis of lateralisation is apparent in different taxa [11,34–39]. In fish and birds [11,32,35,40–42], embryonic light stimulation triggers anatomical and functional lateralisation building on pre-existing endogenous-driven asymmetries. In birds, the rightward torsion of the embryo [42–44] asymmetrically exposes the right eye to the light that filters through the egg, while the left eye is covered by the body (see Figure 1A). This asymmetrical light exposure induces both anatomical and functional lateralisation [19,44–49]. In light-incubated chick embryos, light reaches the right retina (first stage of the input), that feeds the telencephalon (see Figure 1B) via the tectofugal and thalamofugal pathways (see orange and red arrows in Figure 1B for the right-fed pathway). We analysed the effect of light and its effect at different times on the retina and telencephalon, which are the initial and last stations of the visual system affected by light exposure.

**Figure 1.**
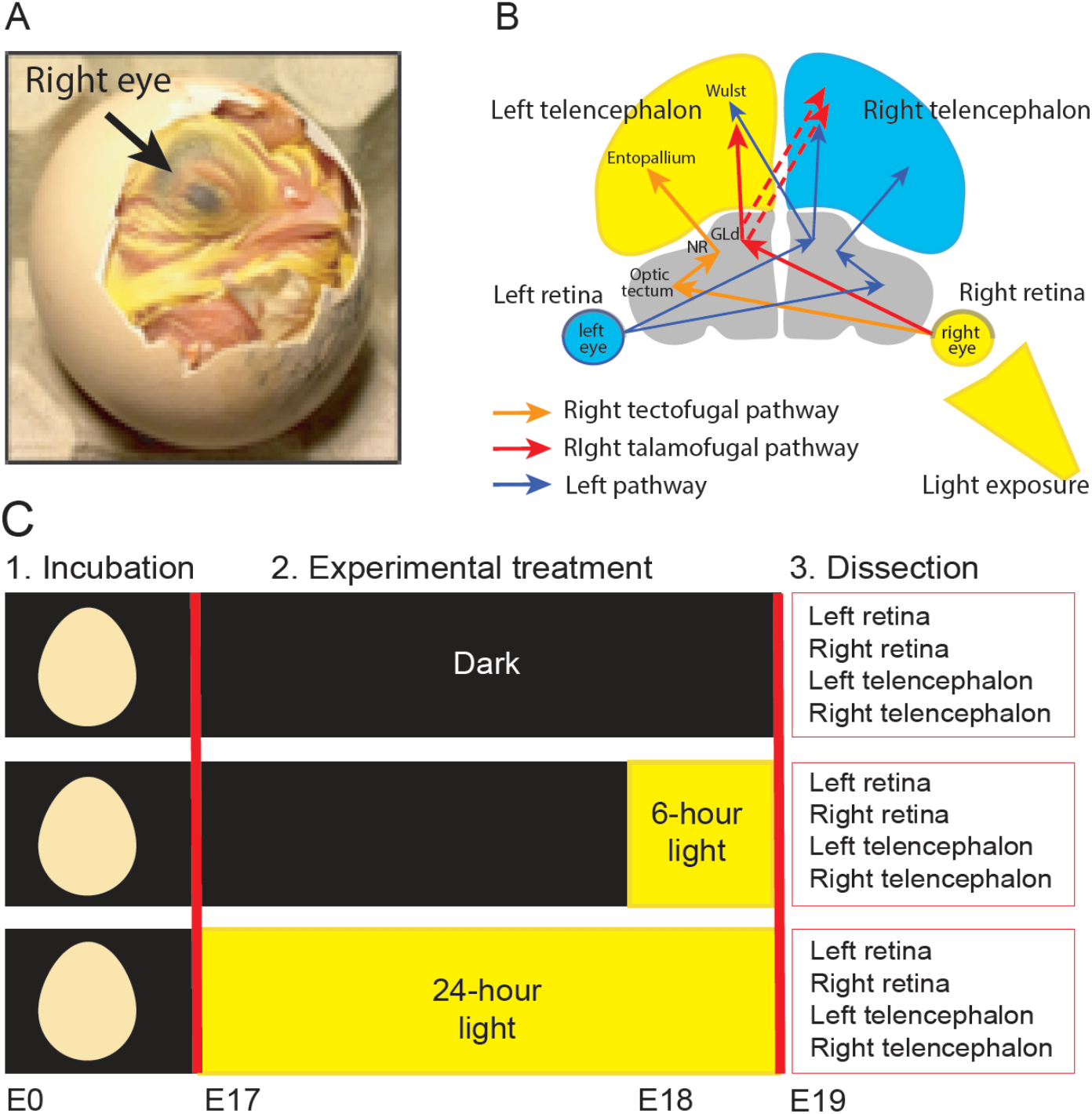
**A. Asymmetric position of chick embryos.** By embryonic day 17 (E17), the right eye is located toward the shell and the left eye tucked under the wing. Only right eye is directly exposed to light. **B. Representation of the visual system in chicks**. The two arrows that depart from the retina indicate the tectofugal and thalamofugal pathways (see [54]). The left tectofugal pathway (orange pathway via right retina, left optic tectum, left nucleus rotundus (NR) and left entopallium in the left telencephalon) and left thalamofugal pathway (red pathway via right retina, left thalamic nucleus geniculatus lateralis pars dorsalis (GLd), left Wulst in the left telencephalon and right Wulst in the right telencephalon) are shown. The right tectofugal and thalamofugal pathways (fed by the left retina) are represented in blue. The regions used in the RNA-seq analysis (retina and telencephalon) are coloured. **C. Schematic representation of the experimental phases**. Eggs were incubated in the dark from embryonic day 0 (E0) to embryonic day 17 (E17). On day E17, eggs were randomly assigned to one of three different experimental treatments. One group continued incubation in darkness (Dark), one group was exposed to 6 hours of light (6-hour light) and one group was exposed to 24 hours of light (24-hour light). At the end of the experimental treatment (E19), dissections were conducted. After dissection, sex was determined by PCR. RNA-seq analysis was carried out on samples from 5 males per light regime.

We focused on the domestic chick (*Gallus gallus*), a prominent model to investigate light-induced lateralisation [40,44,47,50–52]. This model has several advantages: the ease of manipulating light exposure during egg incubation and post-hatch [34,50], the almost complete decussation of the fibres of the optic nerve and reduced number of connections between the two brain hemispheres present in birds [46], and the ease of testing a precocial species that has a mature motor and sensory system at the beginning of life [53].

The effect of embryonic light exposure on lateralisation is apparent at specific time windows [19,34,39,40,44,50]. In chicks, two hours of light exposure in the last 3 days of incubation (E17-E19) are sufficient to determine light-induced behavioural lateralisation, while 6 hours of light exposure make this lateralisation irreversible by the temporary occlusion of the right eye with an eye patch[34]. This transient time window for the induction of environmental-driven lateralisation exhibits the distinctive features of sensitive periods [34,55]. Sensitive periods are specific times during development, in which specific experience-dependent events tune connectivity patterns within the functional range [56,57]. The closure of sensitive periods of brain plasticity can be self-terminated by experience, including exposure to light and visual stimuli, as is clear for filial imprinting in chicks [55,58]. The closure of sensitive periods and neural plasticity are regulated by molecular events that halt/enhance neural plasticity. Many molecular factors that initiate and close plasticity in sensitive periods, from neuromodulatory signals to synaptic proteins and components of the extracellular matrix, are influenced by visual experience [reviewed in 56]. One example is the control of thyroid hormone signalling, mediated by the type-2 deiodinase/iodothyronine deiodinase 2 (DIO2) [59, see 60 for chicks]. Molecules associated with the sensitive periods that regulate the effect of light on lateralisation have not been identified yet.

A broader issue about the ontogenesis of lateralisation is whether environmental exposure to light acts through the same pathway of endogenous asymmetrical mechanisms that produce embryonic rotation and bending towards the right in vertebrate species [61–63], or via a separate mechanism. To shed light on the molecular pathways involved in light-driven lateralisation in chicks, we investigated the effect of light exposure on gene expression in the right and left retinas and telencephalon through an RNA-seq analysis. We exposed E17 embryos to three experimental conditions: complete darkness (Dark), 5-8 hours light exposure (6-hour light), 24-27 hours of light exposure (24-hour light) (Figure 2A). By exposing embryos to at least 5 hours of light, we targeted responses subsequent to those of immediate early genes [64] and reached the amount of stimulation (between 2.5 to 6 hours of light exposure) that provides irreversible lateralisation in chicks [34]. With this design, we looked at the short term and sustained effect of embryonic light exposure in the retina and telencephalon compared to no light exposure.

**Figure 2.**
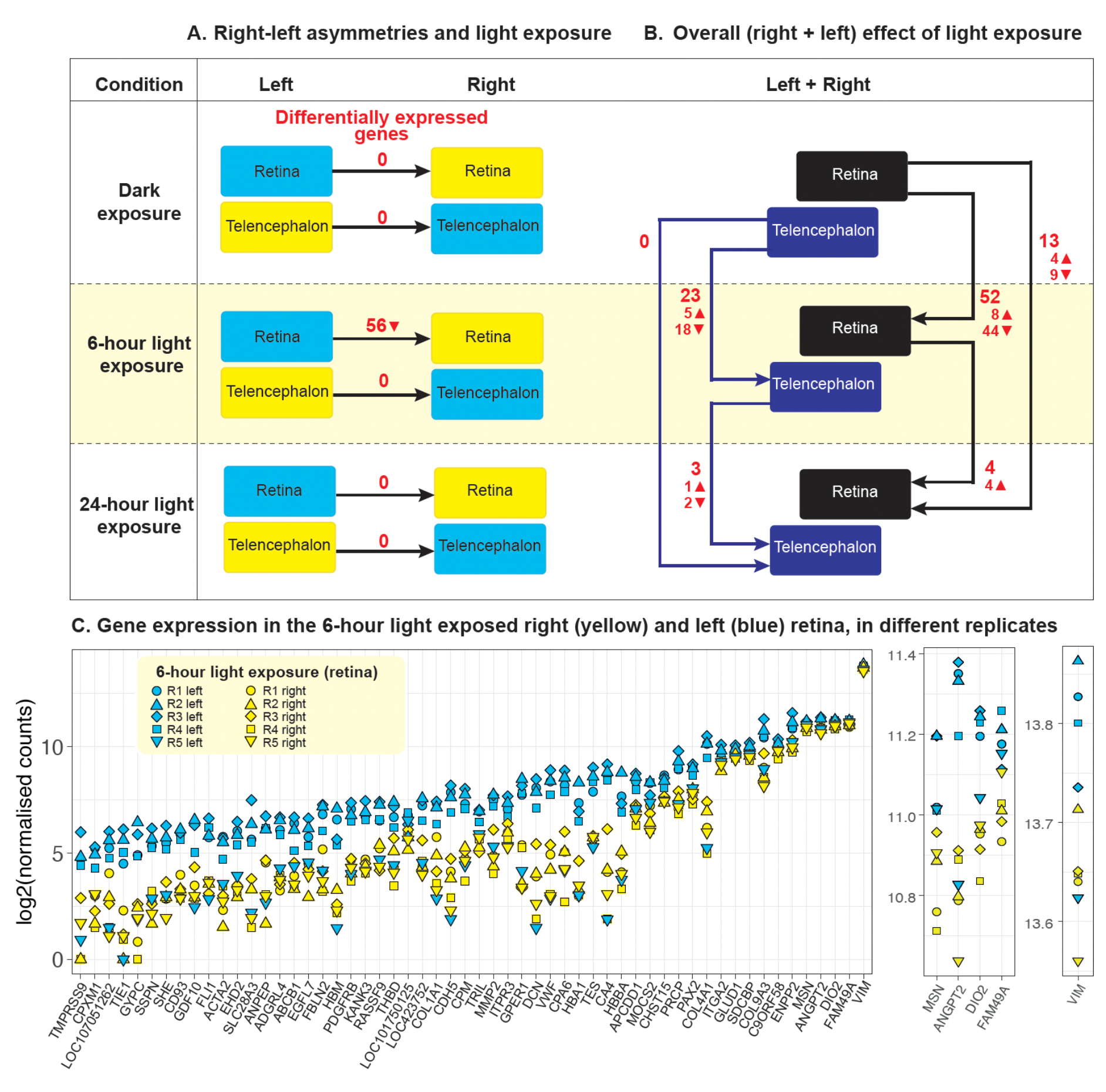
**A. Diagram of the left-right comparisons** and differentially expressed genes (in red) for each comparison (▲upregulation, ▼downregulation) **B. Diagram of the overall (right + left) effect of different exposure to light** (darkness, 6-hour, 24-hour) as differentially expressed genes (in red) in the retina and telencephalon. **C. Gene expression**. For each gene differentially expressed in the retina at 6-hour light exposure, the log2(normalised counts) are shown on the y axis. A supplementary close-up is presented on the right panels for better visualisation of all replicates. Yellow symbols indicate the replicates of the right retina, blue symbols indicate replicates of the left retina. Different symbols indicate different replicas, as shown in the legend.

## RESULTS

### Differential gene expression: lateralised and overall effects

To identify genes differentially expressed in the right and left hemisphere in the absence or presence of asymmetric light exposure, we compared left vs right gene expression in both tissues (retina and telencephalon) in the Dark, 6-hour light and 24-hour light conditions, using a within-subjects design (Figure 2A). In the retina, we identified 56 differentially expressed (DE) genes (adjusted p-value <0.05, Supplementary Table 1) in the right vs left side after 6 hours of light exposure. All differentially expressed genes were downregulated in the right vs. left retina. Besides genes involved in the circulatory system development and extracellular matrix/cell motility (see gene set enrichment analysis below) we identified as differentially expressed: *DIO2*, a gene involved in sensitive periods[59,60], as differentially expressed; *Pax2*, that plays an important role in the glia during retinal development [65]. We found no significant left-right differences in gene expression in either darkness or after 24 hours of light exposure in the retina. No significant differences in gene expression were identified in the telencephalon.

To identify the overall, non-lateralised effect of light exposure, we compared gene expression of both brain sides (i.e., right plus left retina, right plus left telencephalon) between different light exposure conditions (Dark vs 6-hour light, Dark vs 24-hour light, 6-hour light vs 24-hour light) (Figure 2B). In both tissues, the largest effect has been found in the Dark vs 6-hour light exposure comparison (53 differentially expressed genes in the retina, 23 in the telencephalon). In the 6-hour vs 24-hour comparison we found 4 DE genes in the retina, 3 in the telencephalon; in the dark vs 24-hour exposure comparison we found 13 DE genes in the retina (4 upregulated, 9 downregulated, including CAPN15, that is involved in the development of the visual system) and 0 in the telencephalon (see Supplementary Tables 2A-E). The reduction of DE genes after more than 6 hours of light exposure corroborates behavioural and neuroanatomic evidence that, in different species, embryonic light exposure has a transient effect [e.g. see 40 for different time windows in chicks, see 66 for zebrafish]. Behavioural evidence suggests that, in domestic chick embryos, this self-terminating sensitive period ends after 6 hours of light exposure during the last three days of incubation [34].

### Gene set enrichment analysis

To identify biological pathways connected to significant asymmetrical light exposure, we used the differentially expressed proteins identified above (Retina dark vs 6-hour exposure) and performed a gene enrichment analysis via g:Profiler [67,68], with 16356 protein-coding genes used in the model as the background, and FDR at 1%. In the Retina dark vs 6-hour exposure comparison, 32 biological process GO terms, 14 cellular component terms and 23 molecular function GO terms were enriched (see Figure 3 and full enriched terms in Supplementary Table 3).

**Figure 3.**
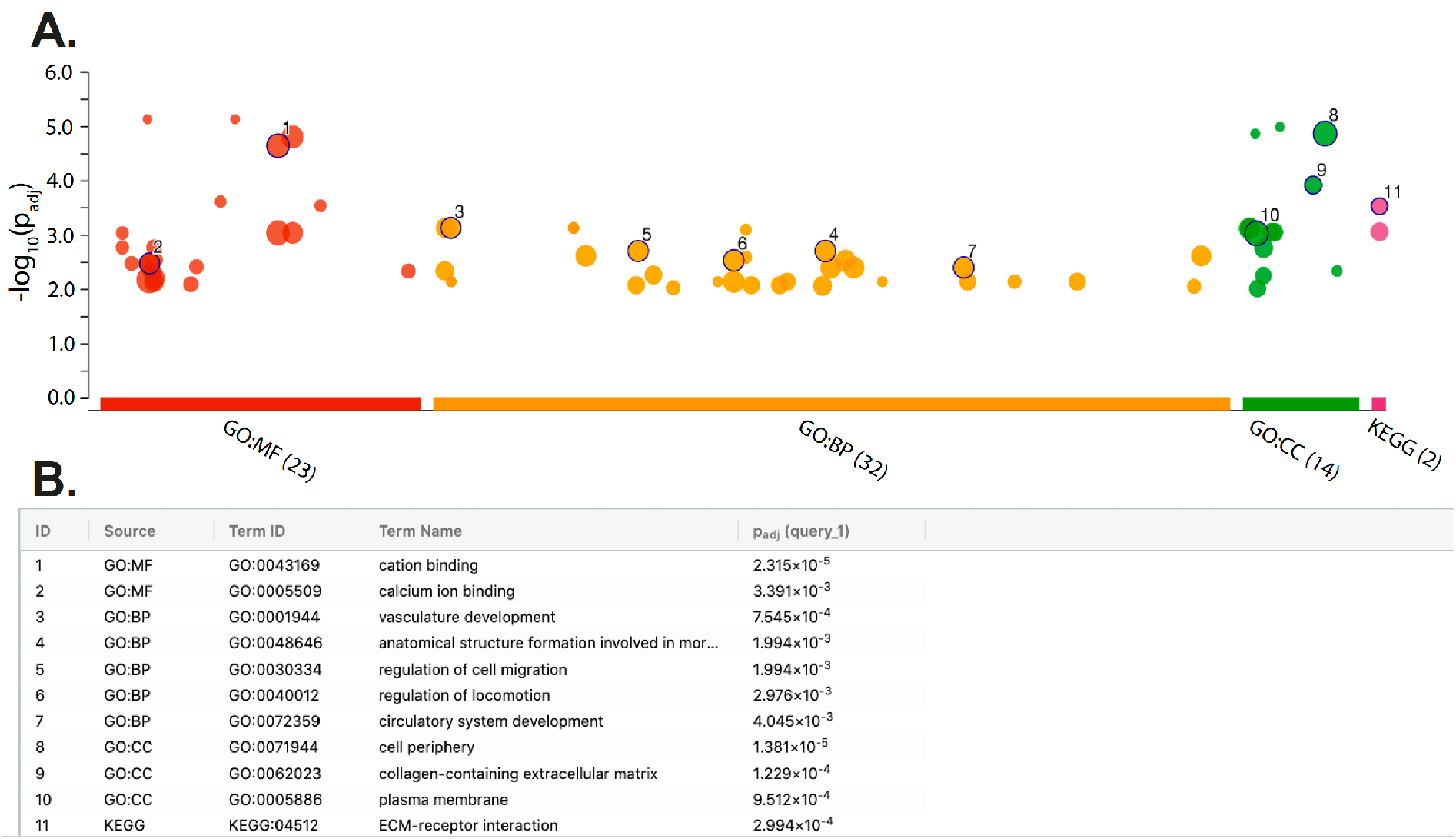
Gene enrichment analysis: functional terms and KEGG pathways. **A. g:Profiler Manhattan plot**. The x axis shows the functional terms (term size greater than 50), grouped and colour-coded by data source (red = GO molecular function, orange = GO biological process, green = GO cellular component, pink = KEGG database). The position of terms in the plots is fixed and terms from the same gene ontology branch are close to each other. The size of circles shows the corresponding term size. Numbers correspond to the terms in the table below; the full list of terms is available in Supplementary Table 3. **B. Legend gene enrichment analysis**. The table shows the ID referred to the Manhattan plot above, source database, term ID, term name, adjusted p values after FDR correction. See Supplementary Table 3 for the full list of terms.

The enriched biological processes relate to two main systems: vascular/circulatory system development and anatomical structure formation. In line with these results, in studying the functional genetics of handedness and lateralisation, Schmitz and colleagues [33] found enrichment on GO terms related to anatomic structure development, including cardiovascular system development, artery development and epithelial tube morphogenesis. The enriched cellular components include the extracellular matrix (that provides physical scaffolding for the cellular constituents and can initiate biochemical and biomechanical cues required for tissue morphogenesis, differentiation and homeostasis), plasma membrane (GO:0005886), haptoglobin-haemoglobin complex (GO:0031838), cell periphery (GO:0071944) and haemoglobin complex (GO:0005833) as top hits. Finally, the enriched molecular functions include metal ion binding (GO:0046872), cation binding (GO:0043169) and platelet-derived growth factor binding (GO:0048407), haptoglobin binding (GO:0031720) and oxygen carrier activity (GO:0005344) and metallocarboxypeptidase activity (GO:0004181) as top hits. Using the same approach, no functional enrichment was found in other tissues and conditions.

KEGG pathway analysis confirmed the significant enrichments for extracellular matrix (ECM) receptor-interaction, that is important in tissue morphogenesis and in the maintenance of tissue and function, and focal adhesion, that is involved in cell motility proliferation and differentiation. Interestingly, the membrane of primary cilia has been associated with receptors for ECM proteins [69,70]. Similarly, in studying the functional genetics involved in handedness with KEGG pathway analysis, Schmitz and colleagues found enrichment for ECM-receptor interaction and focal adhesion. Moreover, they found enrichment in the TGF-beta signalling pathway. This pathway is connected to PDGFRB, which we found to be differentially expressed in right vs left retina at 6 hours of light exposure. This gene, essential for the development of the cardiovascular system, helps in the rearrangement of the actin cytoskeleton and is involved in the primary cilia mechanism [71]. This further evidence points toward the implication of primary cilia in the experience-driven, light-triggered gene pathway of lateralisation initiated in the eye [13].

Overall, gene set enrichment analysis shows an involvement of blood vessels/circulatory system and processes related to morphogenesis in response to 6 hours of light exposure in the retina. Moreover, we detected a temporal-specific effect in the retina present after 6 hours of light stimulation but not after 24 hours. No enrichment was present in other experimental conditions (bilateral tissue sampled at different light-exposure times) and in the telencephalon.

### Interaction network

To investigate the protein interaction network of the differentially regulated genes involved with embryonic light exposure (left-right asymmetry after 6-hour light exposure), we analysed the differentially expressed genes using STRING v11.0 [72] and visualised its output with Cytoscape [73,74]. We analysed the full network (physical and non-physical interactions) with all available interaction sources and medium confidence cut-off (required interaction score ≥ 0.4). Of the 56 differentially expressed genes for Retina 6-hour light exposure, 53 could be found in STRING. The 53 nodes in the overall network produce 40 interactions (vs 6 expected interactions if the nodes were selected at random), with a significant protein-protein interaction enrichment (p<1.0e-16), involving 25 genes. Two interaction subnetworks have been identified (see Fig 4).

**Figure 4.**
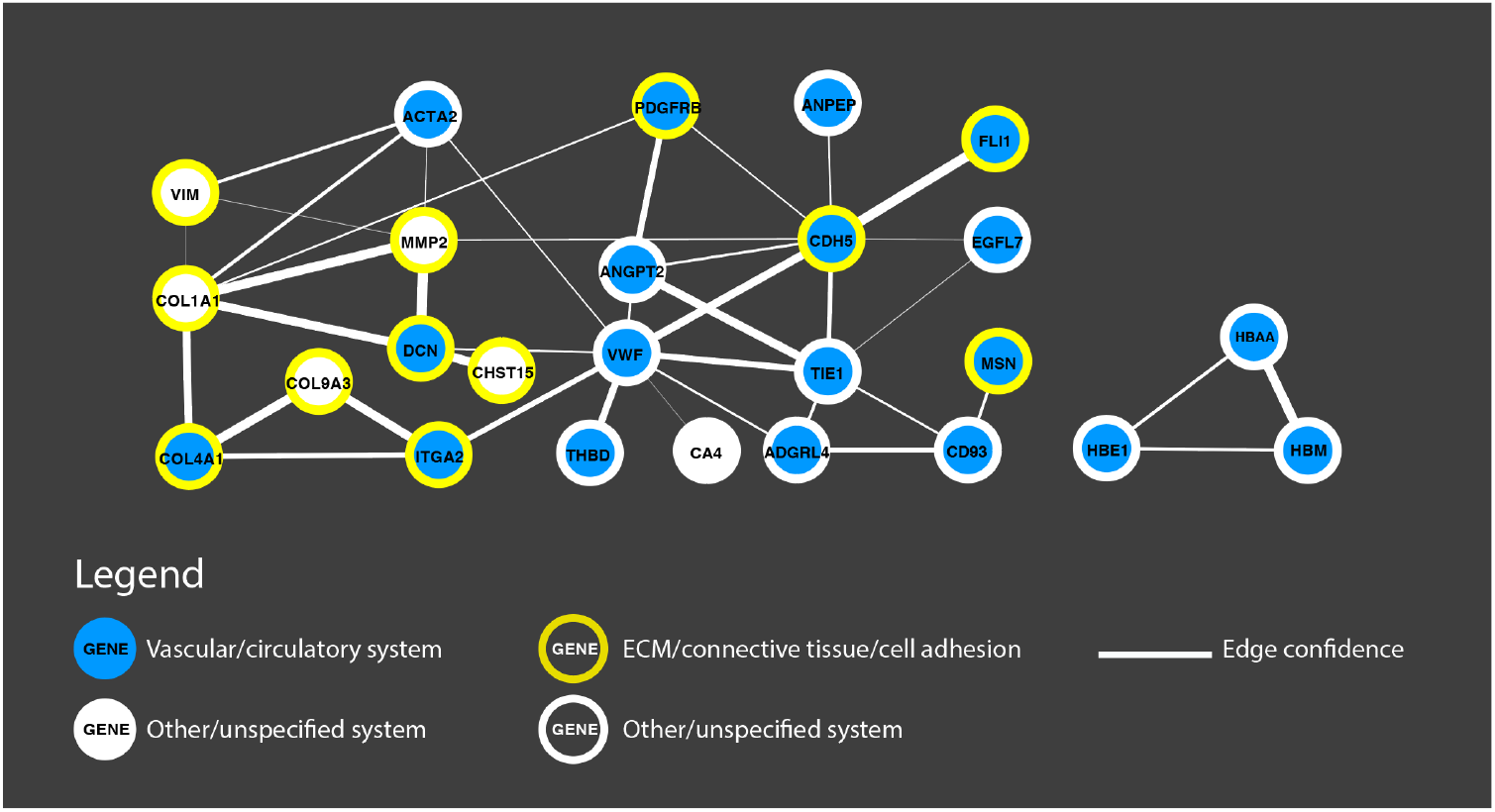
**Interaction network** of genes differentially expressed in the right vs left retina in the 6-hour light exposure condition. Physical and non-physical interactions are represented, based on all available sources in Cytoscape [73] (text mining, experiments, databases, co-expression, neighbours, gene fusion and co-occurrence). Involved systems and functions (e.g., vascular/circulatory system, extracellular matrix (ECM)/connective tissue/cell adhesion are coloured according to the legend, larger edges indicate stronger confidence in the interaction.

The main network (left in Fig. 4) includes nodes involved in the vascular/circulatory system (blue circle) and in connective tissue/cell adhesion/extracellular matrix (yellow outline); the haemoglobin-complex subnetwork (right) is linked to the vascular/circulatory system.

## Discussion

The mechanisms that drive the ontogenesis of lateralisation are far from being understood. We used chicken embryos to identify genetic pathways involved in lateralisation induced by asymmetrical light stimulation. We compared gene expression in the right and left retinas and telencephalon in embryos exposed to different light regimes. In natural settings, light exposure is regulated by the parents, that systematically interrupt incubation in the last days before hatching[75,76]. In these periods, light reaches the egg and embryo.

After exposing embryos to darkness, 6-hour light or 24-hour light, we dissected samples at the same developmental stage (embryonic day E19) and compared the effects of endogenous and stimulus-driven lateralisation induced by light in the right and left brain. We identified significantly differentially expressed genes in the retina in the 6-hour light exposure condition (Figure 2A). Because all samples have been collected at the same developmental stage, we conclude that the observed differential gene expression between left and right retina is due to light exposure.

Our findings support the idea that differential gene expression induced by light exposure terminates between 6 and 24 hours of light exposure in the retina. The lack of differences observed after 24 hours of light exposure while differences are detected at 6 hours of light exposure confirms previous findings about the presence of transient sensitive periods in the development of the nervous system in chicks [for chicks see 55,77,78], and other vertebrates [59,79] (see [80] for developmental changes in human foetuses). Moreover, the timing of this effect is in line with the fact that inversion of lateralisation in chicks has been achieved only between 2.5 and 6 hours of light exposure at E19/E20 [34]. Overall, this pattern of data suggests that, between 6 and 24 hours of light exposure the sensitive period of neural plasticity that stabilises lateralisation in the retina self-terminates. This hypothesis is corroborated by the significant differential expression of DIO2 (iodothyronine deiodinase 2) in the 6-hour light exposure but not in the 24-hour light exposure. DIO2 is a thyroid hormone converting enzyme with an essential role in neurodevelopment and the regulation of the sensitive periods in mammals and birds [59]. Modulation of DIO2 can finely tune the availability of triiodothyronine at specific times during brain development. This gene regulates the start of the sensitive period for filial imprinting in chicks (the self-terminating mechanism that mediates affiliative learning), as well as the regulation of predisposed preference in domestic chicks, such as the preference for biological motion [81,82] and objects that move changing in speed [78]. Evidence that light exposure has a transient effect in a specific time window comes also by the comparison of the samples from both sides of each tissue (right plus left retina, right plus left telencephalon). In these comparisons, in both tissues we identified more differentially expressed genes in the darkness vs 6-hour light exposure than in the 6-hour light exposure vs 24-hour light exposure conditions (23 vs 3 DE genes in the telencephalon; 52 vs 4 DE genes in the retina, see Fig. 2B).

While the retina showed right vs left differential gene expression at 6 hours of light exposure, we did not detect any difference between sides in the telencephalon. A similar lack of significant left-right differences in telencephalon has been previously reported in fish [83] and in human foetuses in the perisylvian area (neocortex), an area with high interhemispheric asymmetry [84], while only mild and age-dependent asymmetries have been recently found in human foetuses[80]).

The telencephalon is a large portion of the chicken pallium fed by both the tectofugal and thalamofugal pathways (see Figure 1B). Lack of lateralised gene expression at E19 can hence be due to different factors. Since we dissected a large portion of the brain, the lack of differential expression could be due to more regionally restricted lateralised effects mediated by light. In addition, light exposure may have its maximum effect in the telencephalon more than 24 hours after the beginning of light stimulation. Moreover, the stimulation of both tectofugal and thalamofugal pathways may activate both hemispheres, since the thalamofugal pathway has both ipsilateral and contralateral projections in the telencephalon. Further experiments should clarify whether more anatomically restricted areas of the embryonic telencephalon are affected by light exposure and on what time course.

Gene enrichment analyses have revealed two main pathways associated with lateralised differences induced by light stimulation: the vascular/circulatory system and the extracellular matrix (ECM). An interesting overlap between gene networks involved in the vascular system and the ECM/connective tissue appears in the interaction network analysis (Figure 4). It is worth discussing the connection between the vascular system, ECM and lateralisation in more detail. As for the vascular system, one of the first macroscopic asymmetries observed during vertebrate development is the right-side looping of the heart tube [see 62 for chicks, reviewed in 85]. More in general, organogenesis is a highly asymmetric process and the biophysical mechanisms engaged in organogenesis involve the ECM. The ECM is engaged in organogenesis via proliferation and traction forces that are mediated for instance by the cytoskeleton [reviewed in 86]. Together with previous literature, our findings suggest a connection between the vascular system and extracellular matrix in establishing asymmetries. Overall, the involvement of blood vessels/vascular system and extracellular matrix in asymmetries support the hypothesis that genes associated with lateralisation might be involved in anatomic structure development guided by environmental cues, rather than in asymmetrical body formation per se (see also [33]).

The ECM is connected to ion activity and to the asymmetric movement of primary cilia, that drives asymmetries [22]. At the molecular level, we found enrichment of ion activity, including Ca^2+^ ion binding, that is connected to the activity of cilia. Previous reports have highlighted the role of calcium ions in lateralisation (e.g. fish [87], mice [26], chicks [88]). The membrane of primary cilia is associated with receptors for ECM proteins [69,70]. Enrichment for ECM-receptor interaction and focal adhesion has been found in the functional genetics of handedness [33], as well as enrichment in the TGF-beta signalling pathway. This pathway is connected to PDGFRB (platelet-derived growth factor receptor β), one of the differentially expressed genes in the 6-hour light exposure left-right comparison. This gene, essential for the development of the cardiovascular system, helps in the rearrangement of the actin cytoskeleton and is involved in the primary cilia mechanism [71]. Overall, our results suggest that there are similarities between the endogenous lateralisation mechanisms and environmentally driven lateralisation since in both cases there is an involvement the extracellular matrix and the platelet-derived growth factor receptor PDGFRB.

Among the differentially expressed gene in the right vs left retina is *Pax2*, a paired like transcription factor involved in the generation of the optic stalk and in the first stage of chiasm asymmetric axonal migration [65,89]. *Pax2* expression has also been reported in the chick retina at later developmental stages in Müller glial cells, astrocyte-like radial cells that can undergo reprogramming and repair in teleost fish and regenerate retinal neurons through a process of dedifferentiation and acquisition of multipotent progenitor identity [90, see also 91 for a review].

To summarise, our data show a molecular connection between endogenous and experience-driven lateralisation mediated by light exposure. We identified functional gene networks connected to the vascular system and extracellular matrix as mediators of light-induced lateralisation in the retina, and their connections. The evidence of a temporally defined window of these effects calls for further studies on modulating lateralisation via the molecular pathways that open and close sensitive periods in the developing brain, such as thyroid hormones [59,78,92], with important biomedical implications.

## Supporting information

Overview_all_supplementaryTables

## Availability of Data

All of the sequencing data used are available in the NCBI Short Read Archive under the Bioproject PRJNA743180 All code used in this analysis is available online at https://github.com/peterthorpe5/Gallusggal6_RNAseq.

## Funding

This project was funded by BBSRC grant BB/S003223/1 (EV, JG), Royal Society grant UF150663 (SP). The funders had no role in study design, data collection and analysis, decision to publish, or preparation of the manuscript.

## Supplementary Methods

### Samples and study design

Sixty-three freshly fertilized eggs of Ross 308 (Aviagen) strain were stored in darkness at 4°C until the beginning of incubation. Eggs were incubated at a constant temperature of 37.7°C and 40% humidity and rotated automatically in complete darkness from embryonic day 0 (E0) to embryonic day 17 (E17) at 9:00 am. Subsequently, eggs were randomly assigned to experimental groups: in the Dark condition eggs continued to stay in darkness, eggs in the 24-hour light exposure condition received light from E17 9:00 am to ~E18 12:00 pm (i.e., 24-27 consecutive hours of light), and eggs in the 6-hour light exposure condition received light from E18 4:00 am to ~E18 12:00 pm (i.e., 5-8 consecutive hours of light) (Figure 1A). This timing of exposure and dissection was chosen to make sure light exposure was long enough to induce functional lateralisation [50] and at the same time target responses subsequent to those of immediate early genes [64]. Light was delivered to eggs through a 5 white-light LEDs of (18 lumens per LED) located at 24 cm from the eggs. Dissections were conducted on E18 between 9 am-12 pm.

### Dissection

A total of 55 embryos were dissected. For each embryo, the egg was retrieved from the incubator, placed at −20°C for 2 min to prevent major blood loss during dissection and then immediately opened by cutting along the widest part of the shell with scissors. Embryonic membranes were then disrupted and the embryo was culled by rapid decapitation followed by the removal of both eyes and then brain. The dissection of retinas and telencephalons were performed in parallel by two experimenters to minimize sample collection time. Brain tissue was macroscopically subdivided in two regions: telencephalon and remainder of the brain. Of these regions, only telencephalon was further subdivided into left and right samples and processed for RNA-seq. Macrodissected brain samples were frozen immediately on dry ice and stored at −80°C until RNA extraction. For retina dissection, first, eyelids were cut away and the whole eye was carefully removed without piercing the eye globe. Then, dissection of the retina from the enucleated eyes was performed by cutting the eyes in 1⁄2 along their equator exposing the vitreous and retina. The translucent gelatinous vitreous humor was carefully removed with forceps exposing the retina. Retina and pigmented epithelium were removed from the eye cup and placed immediately on dry ice and stored at −80°C until RNA extraction.

### Molecular sexing

Since anatomic asymmetries in the thalamofugal pathway [93] and behavioural asymmetries [94,95] are more pronounced in male chicks compared to females [reviewed in 96], we focused our study on males only.

### RNA extraction and quality control

Total RNA was extracted and DNAse-treated using RNeasy mini kit (Qiagen) following the manufacturer’s protocol. RNA concentrations were measured using the Qubit RNA Broad-Range Assay kit and RNA quality was analysed on an Agilent Bioanalyzer 2100. Only samples with RNA integrity numbers (RIN) ≥ 9 were considered for RNA-seq inclusion.

### RNA-seq analysis

RNA-seq data were generated from a total of 60 samples on an Illumina HiSeq 4000 at the Genomic facilities of the Wellcome Trust Human Genetic Centre (WTHGC), University of Oxford. Five biological replicates were used for each tissue and condition: dark, 6-hour and 24-hour light exposure for the retina and telencephalon on both the left and right side. One μg of input RNA was used for the standard TruSeq Stranded mRNA Library Preparation Kit that includes isolation of mRNA by polyA-selection. The libraries were then pooled according to their tissue, side and treatment group. The sequence libraries had a median number of 11,232,013 reads before trimming (10261901 reads after trimming).

All data were checked for quality with FastQC (FastQC v0.11.5, 2017 https://www.bioinformatics.babraham.ac.uk/projects/fastqc/) and contaminants with FastQ Screen (v0.10.0), then trimmed for adapter sequences with Trimmomatic (v0.36) [97]. The statistics of number of reads per sample are shown in Supplementary Table 4.

### Differential gene expression analysis

Surviving reads were quantified by mapping against the Ensembl galGal6 chicken transcriptome (downloaded from ftp://ftp.ncbi.nlm.nih.gov/genomes/all/GCF/000/002/315/GCF_000002315.6_GRCg6a/GCF_000002315.6_GRCg6a_rna.fna.gz) using Salmon (v0.12.0) [98]. Mapping efficiency ranged from 76-81%. Transcript read counts were collapsed to genes with tximport (ref) in R version 4.0.3 (https://www.R-project/org), then counts and fragment lengths were imported into DESeq2 (1.30.0)) [99]. Principal components analysis (PCA) was applied to visualize variance in the data, and one individual in the 24-hour light exposure treatment group was excluded from further analysis because it was an outlier and it was annotated as underdeveloped at the time of dissection. Sequencing replicates of each sample library were similar by PCA, so they were combined to increase statistical power. Differential gene expression analysis was conducted separately for retina and telencephalon tissues. With DESeq2, we compared gene expression in left and right sides of the same tissue in the same treatment conditions, adding a factor to our model to control for individual effects, and we also looked for aggregate differences in gene expression between treatment groups (reference code in Available data for details). An adjusted p-value cut-off of p<0.05 was applied for all comparisons. Analysis was conducted with R (https://www.R-project.org/) and RStudio (http://www.rstudio.com). The packages dplyR (https://CRAN.R-project.org/package=dplyr), biomaRt [100], tximport [101] and independent filtering was employed to optimize detection of genes below this threshold. Shrinkage estimation was applied to moderate fold-changes of low abundance genes to facilitate ranking for downstream analysis of gene ontology.

## Notes

### Competing Interest Statement

The authors have declared no competing interest.

