## Supplementary material for "Light-induced asymmetries in the embryonic retina are mediated by the vascular system and extracellular matrix": Overview_all_supplementaryTables

### **Supplementary Tables** (see also individual spreadsheets)

#### Supplementary table 1

Differentially expressed genes in the left-right comparison (adjusted p-value p<0.05, left considered as baseline).

| tissue | condition | comparison | gene | baseMean | log2FoldChange | lfcSE | pvalue | padj |
|---|---|---|---|---|---|---|---|---|
| retina | 6-hour exposure | left-right | HBM | 50.1580938 | -2.8617565 | 0.60189424 | 1.57E-08 | 0.00025669 |
| retina | 6-hour exposure | left-right | APCDD1 | 180.206098 | -1.1681799 | 0.25847546 | 1.88E-07 | 0.00133795 |
| retina | 6-hour exposure | left-right | SLC28A3 | 29.0823833 | -6.59E-06 | 0.0014427 | 2.45E-07 | 0.00133795 |
| retina | 6-hour exposure | left-right | ANGPT2 | 2022.84198 | -0.3731778 | 0.0881928 | 8.83E-07 | 0.00196894 |
| retina | 6-hour exposure | left-right | COL1A1 | 70.2769096 | -1.834667 | 0.46439475 | 1.00E-06 | 0.00196894 |
| retina | 6-hour exposure | left-right | LOC423752 | 63.8093406 | -1.9292547 | 0.4695861 | 6.53E-07 | 0.00196894 |
| retina | 6-hour exposure | left-right | MOCS2 | 218.877264 | -1.3391908 | 0.32553767 | 1.08E-06 | 0.00196894 |
| retina | 6-hour exposure | left-right | PDGFRB | 50.4838055 | -1.9367839 | 0.48891916 | 7.87E-07 | 0.00196894 |
| retina | 6-hour exposure | left-right | SSPN | 18.8887878 | -2.475247 | 0.5868452 | 5.71E-07 | 0.00196894 |
| retina | 6-hour exposure | left-right | ADGRL4 | 33.5598194 | -2.0739368 | 0.53205476 | 1.57E-06 | 0.00257034 |
| retina | 6-hour exposure | left-right | FBLN2 | 44.516035 | -2.1652072 | 0.57596597 | 1.74E-06 | 0.0025816 |
| retina | 6-hour exposure | left-right | KANK3 | 52.7021535 | -1.8670177 | 0.49166613 | 2.07E-06 | 0.00282594 |
| retina | 6-hour exposure | left-right | HBBA | 152.248634 | -5.64E-06 | 0.0014427 | 3.14E-06 | 0.00394851 |
| retina | 6-hour exposure | left-right | HBA1 | 116.585849 | -4.31E-06 | 0.0014427 | 4.51E-06 | 0.00525568 |
| retina | 6-hour exposure | left-right | SHE | 20.5263901 | -7.98E-06 | 0.0014427 | 4.82E-06 | 0.00525568 |
| retina | 6-hour exposure | left-right | CDH5 | 70.7658541 | -4.16E-06 | 0.0014427 | 8.59E-06 | 0.00877687 |
| retina | 6-hour exposure | left-right | SDCBP | 806.800793 | -0.3522846 | 0.09567858 | 1.04E-05 | 0.00943533 |
| retina | 6-hour exposure | left-right | VWF | 110.928778 | -3.72E-06 | 0.0014427 | 1.04E-05 | 0.00943533 |
| retina | 6-hour exposure | left-right | ACTA2 | 28.4215603 | -2.2040505 | 0.66022928 | 1.27E-05 | 0.00992613 |
| retina | 6-hour exposure | left-right | COL4A1 | 391.785186 | -3.97E-06 | 0.0014427 | 1.22E-05 | 0.00992613 |
| retina | 6-hour exposure | left-right | EHD2 | 28.6211121 | -2.1067567 | 0.61211054 | 1.19E-05 | 0.00992613 |
| retina | 6-hour exposure | left-right | GDF10 | 26.7562964 | -1.8887349 | 0.57799457 | 1.40E-05 | 0.01039492 |
| retina | 6-hour exposure | left-right | GYPC | 16.4306326 | -3.246276 | 1.08675434 | 1.48E-05 | 0.01055033 |
| retina | 6-hour exposure | left-right | EGFL7 | 35.0878888 | -1.3728233 | 0.40851093 | 1.75E-05 | 0.01177886 |
| retina | 6-hour exposure | left-right | MMP2 | 78.0528065 | -6.61E-06 | 0.0014427 | 1.85E-05 | 0.01177886 |
| retina | 6-hour exposure | left-right | TIE1 | 15.9303226 | -3.1841369 | 1.05059633 | 1.87E-05 | 0.01177886 |
| retina | 6-hour exposure | left-right | GPER1 | 80.8276903 | -2.6307098 | 0.94617532 | 2.04E-05 | 0.01235911 |
| retina | 6-hour exposure | left-right | FLI1 | 27.1188037 | -1.8930093 | 0.63456381 | 2.47E-05 | 0.01398514 |
| retina | 6-hour exposure | left-right | MSN | 1916.37018 | -0.2085911 | 0.06157933 | 2.48E-05 | 0.01398514 |
| retina | 6-hour exposure | left-right | TRIL | 72.2526573 | -1.52E-05 | 0.00144272 | 2.60E-05 | 0.01416624 |
| retina | 6-hour exposure | left-right | CHST15 | 230.27279 | -0.5943206 | 0.18320287 | 3.56E-05 | 0.0187957 |
| retina | 6-hour exposure | left-right | DIO2 | 2078.68668 | -0.225633 | 0.07091721 | 4.93E-05 | 0.02445721 |
| retina | 6-hour exposure | left-right | VIM | 13248.9161 | -0.1093278 | 0.03427688 | 4.85E-05 | 0.02445721 |
| retina | 6-hour exposure | left-right | THBD | 54.5328539 | -1.6046616 | 0.53504641 | 5.19E-05 | 0.0249628 |
| retina | 6-hour exposure | left-right | ABCB1 | 33.6414107 | -7.46E-06 | 0.0014427 | 5.46E-05 | 0.02550097 |
| retina | 6-hour exposure | left-right | DCN | 89.5467718 | -2.6922775 | 1.21231532 | 6.02E-05 | 0.02734692 |

| | | | | | | | | |
|---|---|---|---|---|---|---|---|---|
| retina | 6-hour exposure | left-right | C9ORF58 | 916.215407 | -4.56E-05 | 0.00144288 | 6.41E-05 | 0.02835856 |
| retina | 6-hour exposure | left-right | CPM | 70.8064595 | -5.18E-06 | 0.0014427 | 7.39E-05 | 0.03129629 |
| retina | 6-hour exposure | left-right | ENPP2 | 1345.77197 | -0.8251012 | 0.27443552 | 7.46E-05 | 0.03129629 |
| retina | 6-hour exposure | left-right | ANPEP | 30.7450329 | -6.07E-06 | 0.0014427 | 8.41E-05 | 0.03355182 |
| retina | 6-hour exposure | left-right | RASSF9 | 53.2819187 | -7.53E-06 | 0.0014427 | 8.35E-05 | 0.03355182 |
| retina | 6-hour exposure | left-right | COL9A3 | 821.570351 | -1.3404996 | 0.45968154 | 8.83E-05 | 0.0343916 |
| retina | 6-hour exposure | left-right | CPA6 | 111.877333 | -3.35E-06 | 0.0014427 | 9.78E-05 | 0.03549686 |
| retina | 6-hour exposure | left-right | CPXM1 | 13.6240995 | -1.9618093 | 0.72015794 | 9.77E-05 | 0.03549686 |
| retina | 6-hour exposure | left-right | FAM49A | 2130.15677 | -9.48E-05 | 0.00144344 | 9.98E-05 | 0.03549686 |
| retina | 6-hour exposure | left-right | ITGA2 | 661.340654 | -0.4326308 | 0.14652649 | 9.40E-05 | 0.03549686 |
| retina | 6-hour exposure | left-right | CA4 | 145.642959 | -1.94E-06 | 0.0014427 | 0.00010497 | 0.03577066 |
| retina | 6-hour exposure | left-right | TMPRSS9 | 9.17652102 | -3.2468441 | 1.63939989 | 0.00010414 | 0.03577066 |
| retina | 6-hour exposure | left-right | ITPR3 | 80.3484197 | -0.8371948 | 0.29165194 | 0.00011322 | 0.03779527 |
| retina | 6-hour exposure | left-right | PAX2 | 315.213701 | -0.918332 | 0.32432951 | 0.00011706 | 0.03829432 |
| retina | 6-hour exposure | left-right | PRCP | 286.897413 | -8.48E-06 | 0.0014427 | 0.00012135 | 0.03892156 |
| retina | 6-hour exposure | left-right | CD93 | 22.647214 | -1.7754099 | 0.66250772 | 0.00012555 | 0.03949379 |
| retina | 6-hour exposure | left-right | TES | 127.282445 | -5.09E-06 | 0.0014427 | 0.00013056 | 0.04029228 |
| retina | 6-hour exposure | left-right | GLUD1 | 784.236956 | -0.259483 | 0.09077447 | 0.00013338 | 0.04040174 |
| retina | 6-hour exposure | left-right | LOC107051262 | 14.8141011 | -5.15E-06 | 0.0014427 | 0.00016455 | 0.04893574 |
| retina | 6-hour exposure | left-right | LOC101750125 | 62.1671502 | -0.6330772 | 0.22816151 | 0.00016839 | 0.04918393 |

**Supplementary table 2A:** Differentially expressed genes (adjusted p-value p<0.05) in the 24-hour vs 6-hour light condition in the retina (6-hour light used as baseline).

| tissue | condition | comparison | gene | baseMean | log2FoldChange | lfcSE | pvalue | padj |
|---|---|---|---|---|---|---|---|---|
| retina | (left+right) | 24-hour vs 6-hour | GDF3 | 3444.79121 | 0.14193329 | 0.02391898 | 3.06E-09 | 5.93E-05 |
| retina | (left+right) | 24-hour vs 6-hour | DPYSL4 | 3475.65935 | 0.16577801 | 0.03274498 | 4.27E-07 | 0.00413343 |
| retina | (left+right) | 24-hour vs 6-hour | CYTL1 | 7.90748155 | 0.03664545 | 0.00967095 | 1.01E-05 | 0.04877442 |
| retina | (left+right) | 24-hour vs 6-hour | MDGA1 | 2736.60857 | 0.15016454 | 0.03371088 | 8.45E-06 | 0.04877442 |

**Supplementary table 2B:** Differentially expressed genes (adjusted p-value of p<0.05) in the 24-hour light vs dark condition in the retina (dark used as baseline).

.

| tissue | condition | comparison | gene | baseMean | log2FoldChange | lfcSE | pvalue | padj |
|---|---|---|---|---|---|---|---|---|
| retina | (left+right) | 24-hour vs dark | CAPN15 | 1289.28846 | -0.1409149 | 0.02968143 | 9.24E-08 | 0.00144212 |
| retina | (left+right) | 24-hour vs dark | LOC107054696 | 29.3397509 | 0.94444452 | 0.22139474 | 7.67E-07 | 0.0059878 |
| retina | (left+right) | 24-hour vs dark | LOC112530945 | 194.391685 | 1.2556644 | 0.30202571 | 1.38E-06 | 0.00715974 |
| retina | (left+right) | 24-hour vs dark | AMBRA1 | 696.129829 | -0.1400366 | 0.03423815 | 1.90E-06 | 0.0074254 |
| retina | (left+right) | 24-hour vs dark | BZFP2 | 366.475355 | -0.4167702 | 0.11145005 | 6.57E-06 | 0.01704123 |
| retina | (left+right) | 24-hour vs dark | CHD5 | 2487.07944 | -0.1031858 | 0.0268526 | 5.57E-06 | 0.01704123 |
| retina | (left+right) | 24-hour vs dark | LOC107053028 | 12.0100784 | -1.3001211 | 0.3459291 | 7.64E-06 | 0.01704123 |
| retina | (left+right) | 24-hour vs dark | SELENON | 1714.85191 | 0.10305452 | 0.02884806 | 1.56E-05 | 0.03034793 |
| retina | (left+right) | 24-hour vs dark | APOOL | 1473.04098 | 0.18780297 | 0.05597663 | 3.01E-05 | 0.03916261 |
| retina | (left+right) | 24-hour vs dark | CYP2J21 | 939.857032 | -0.3070443 | 0.09148302 | 2.74E-05 | 0.03916261 |
| retina | (left+right) | 24-hour vs dark | DTX3L | 42.0258214 | -0.4939051 | 0.14752738 | 2.76E-05 | 0.03916261 |
| retina | (left+right) | 24-hour vs dark | TOB2 | 1242.76892 | -0.1434636 | 0.04236753 | 2.78E-05 | 0.03916261 |
| retina | (left+right) | 24-hour vs dark | LOC112532332 | 22.0611046 | -0.7249746 | 0.22098272 | 4.09E-05 | 0.04911673 |

**Supplementary table 2C:** Differentially expressed genes (adjusted p-value of p<0.05) in the 6-hour light vs dark condition in the retina (dark used as baseline).

| tissue | condition | comparison | gene | baseMean | log2FoldChange | lfcSE | pvalue | padj |
|---|---|---|---|---|---|---|---|---|
| retina | (left+right) | 6-hour vs dark | MAF1 | 4493.83458 | -0.137085 | 0.02103501 | 3.93E-12 | 2.15E-08 |
| retina | (left+right) | 6-hour vs dark | NPTXR | 4663.3395 | -0.1083198 | 0.02186393 | 2.78E-08 | 7.61E-05 |
| retina | (left+right) | 6-hour vs dark | CHD5 | 2487.07944 | -0.119105 | 0.02543528 | 1.51E-07 | 0.00027588 |
| retina | (left+right) | 6-hour vs dark | CAPN15 | 1289.28846 | -0.1227942 | 0.02913173 | 8.21E-07 | 0.00112325 |
| retina | (left+right) | 6-hour vs dark | GDF3 | 3444.79121 | -0.1034708 | 0.02579164 | 2.61E-06 | 0.00239451 |
| retina | (left+right) | 6-hour vs dark | NPC1 | 2169.34459 | 0.09139384 | 0.0223407 | 2.63E-06 | 0.00239451 |
| retina | (left+right) | 6-hour vs dark | NFIC | 739.921512 | -0.1608188 | 0.04290515 | 7.89E-06 | 0.00539448 |

| | | | | | | | | |
|---|---|---|---|---|---|---|---|---|
| retina | (left+right) | 6-hour vs dark | TIMP2 | 2752.26281 | -0.123161 | 0.03253792 | 7.19E-06 | 0.00539448 |
| retina | (left+right) | 6-hour vs dark | ASIC4 | 2988.34594 | -0.1467267 | 0.042278 | 2.21E-05 | 0.01343779 |
| retina | (left+right) | 6-hour vs dark | CADM3 | 15218.0061 | -0.082878 | 0.0243599 | 3.78E-05 | 0.01678297 |
| retina | (left+right) | 6-hour vs dark | GOT2 | 3094.7356 | -0.0861882 | 0.02517004 | 4.21E-05 | 0.01678297 |
| retina | (left+right) | 6-hour vs dark | LOC107050378 | 1842.72988 | -0.2350084 | 0.0724055 | 4.29E-05 | 0.01678297 |
| retina | (left+right) | 6-hour vs dark | TTC8 | 3067.55794 | 0.09199942 | 0.02709426 | 3.94E-05 | 0.01678297 |
| retina | (left+right) | 6-hour vs dark | WBP2 | 3403.51367 | -0.0824237 | 0.0241507 | 4.10E-05 | 0.01678297 |
| retina | (left+right) | 6-hour vs dark | PPIA | 6670.99149 | -0.1163877 | 0.03597198 | 5.51E-05 | 0.01885377 |
| retina | (left+right) | 6-hour vs dark | RPUSD1 | 1356.13605 | -0.0918137 | 0.02837218 | 5.41E-05 | 0.01885377 |
| retina | (left+right) | 6-hour vs dark | NACAD | 4248.23078 | -0.0800158 | 0.02448693 | 6.67E-05 | 0.02096815 |
| retina | (left+right) | 6-hour vs dark | NTM | 2544.41562 | -0.1093179 | 0.03439634 | 6.90E-05 | 0.02096815 |
| retina | (left+right) | 6-hour vs dark | CSDC2 | 2281.76596 | -0.1090795 | 0.0348777 | 7.75E-05 | 0.02119313 |
| retina | (left+right) | 6-hour vs dark | USP19 | 1595.39798 | -0.080692 | 0.02478922 | 7.57E-05 | 0.02119313 |
| retina | (left+right) | 6-hour vs dark | SLC25A11 | 1190.74281 | -0.1390247 | 0.04727553 | 8.90E-05 | 0.02221212 |
| retina | (left+right) | 6-hour vs dark | TRIM27 | 1177.08347 | -0.1395176 | 0.04532119 | 8.93E-05 | 0.02221212 |
| retina | (left+right) | 6-hour vs dark | DPYSL2 | 24606.7275 | -0.0883114 | 0.02763284 | 0.00010395 | 0.02473093 |
| retina | (left+right) | 6-hour vs dark | C5H11orf58 | 1812.72115 | 0.085387 | 0.02752916 | 0.00011472 | 0.02596168 |
| retina | (left+right) | 6-hour vs dark | CDKN1A | 1596.0125 | -0.1242896 | 0.04176183 | 0.00012242 | 0.02596168 |
| retina | (left+right) | 6-hour vs dark | CPLX2 | 1287.2288 | -0.1505372 | 0.0503278 | 0.00012336 | 0.02596168 |
| retina | (left+right) | 6-hour vs dark | KIF1A | 11786.5516 | -0.0994656 | 0.02939977 | 0.00012912 | 0.02616743 |
| retina | (left+right) | 6-hour vs dark | ACTC2L | 16796.5066 | -0.1878808 | 0.06350407 | 0.00014844 | 0.02707591 |
| retina | (left+right) | 6-hour vs dark | MTSS1L | 3615.84194 | -0.1129803 | 0.03862113 | 0.00014764 | 0.02707591 |
| retina | (left+right) | 6-hour vs dark | RNASEK | 2408.42201 | 0.25218597 | 0.08833669 | 0.00014719 | 0.02707591 |
| retina | (left+right) | 6-hour vs dark | MTSS1 | 8847.73578 | -0.0528928 | 0.01633129 | 0.00015426 | 0.02723008 |
| retina | (left+right) | 6-hour vs dark | MLLT6 | 2386.51526 | -0.1023824 | 0.03536563 | 0.00016747 | 0.02863782 |
| retina | (left+right) | 6-hour vs dark | SNCB | 9825.96925 | -0.1038275 | 0.0361552 | 0.00018265 | 0.03028656 |
| retina | (left+right) | 6-hour vs dark | IMPA1 | 1410.36326 | 0.09391381 | 0.03264535 | 0.00021464 | 0.03454502 |
| retina | (left+right) | 6-hour vs dark | RBFOX2 | 3106.2141 | -0.1019188 | 0.03621501 | 0.00023832 | 0.03622451 |
| retina | (left+right) | 6-hour vs dark | SMARCD2 | 1492.83112 | -0.1277901 | 0.0461663 | 0.00023371 | 0.03622451 |
| retina | (left+right) | 6-hour vs dark | ZFP91 | 2481.64833 | -0.0782778 | 0.02736812 | 0.00025823 | 0.03819072 |
| retina | (left+right) | 6-hour vs dark | FIBCD1 | 5036.34488 | -0.071227 | 0.02458562 | 0.00027527 | 0.03963918 |
| retina | (left+right) | 6-hour vs dark | LOC101750908 | 945.225036 | -0.1059258 | 0.03952039 | 0.0003029 | 0.04249964 |
| retina | (left+right) | 6-hour vs dark | LOC101749377 | 2033.00223 | -0.0692775 | 0.02501958 | 0.00034352 | 0.04699335 |
| retina | (left+right) | 6-hour vs dark | ARHGEF11 | 4313.49981 | -0.0699082 | 0.02431796 | 0.00040357 | 0.04742089 |

| | | | | | | | | |
|---|---|---|---|---|---|---|---|---|
| retina | (left+right) | 6-hour vs dark | CBX6 | 3314.80721 | -0.0974132 | 0.0381058 | 0.00036869 | 0.04742089 |
| retina | (left+right) | 6-hour vs dark | PPP1R7 | 1789.58059 | 0.07245452 | 0.02646218 | 0.0004013 | 0.04742089 |
| retina | (left+right) | 6-hour vs dark | RCN2 | 1672.35267 | 0.09281993 | 0.03497512 | 0.00038938 | 0.04742089 |
| retina | (left+right) | 6-hour vs dark | RPLP1 | 2191.08684 | -0.1334699 | 0.05246735 | 0.00040731 | 0.04742089 |
| retina | (left+right) | 6-hour vs dark | SLC25A1 | 801.773643 | -0.0943005 | 0.03748468 | 0.0003985 | 0.04742089 |
| retina | (left+right) | 6-hour vs dark | STK26 | 2008.06909 | 0.07673962 | 0.02815506 | 0.00039409 | 0.04742089 |
| retina | (left+right) | 6-hour vs dark | EVI5L | 2625.17697 | -0.1022121 | 0.03871843 | 0.00042303 | 0.04822524 |
| retina | (left+right) | 6-hour vs dark | LOC100859276 | 984.67133 | -0.1352874 | 0.05373905 | 0.00044623 | 0.04916069 |
| retina | (left+right) | 6-hour vs dark | TMEM132E | 2815.74628 | -0.1009028 | 0.0386159 | 0.0004492 | 0.04916069 |
| retina | (left+right) | 6-hour vs dark | AHCYL2 | 6022.12161 | -0.0953885 | 0.0374007 | 0.00047127 | 0.04959233 |
| retina | (left+right) | 6-hour vs dark | PLEKHA6 | 2315.16606 | -0.0970003 | 0.03787424 | 0.00046412 | 0.04959233 |

**Supplementary table 2D:** Differentially expressed genes (adjusted p-value of p<0.05) in the 24-hour vs 6-hour light condition in the telencephalon (6-hour light used as baseline).

| **tissue** | **condition** | **comparison** | **gene** | **baseMean** | **log2FoldChange** | **lfcSE** | **pvalue** | **padj** |
|---|---|---|---|---|---|---|---|---|
| telencephalon | (left+right) | 24-hour vs 6-hour | RHOF | 295.380149 | -0.3452298 | 0.05724535 | 1.61E-09 | 0.0000315 |
| telencephalon | (left+right) | 24-hour vs 6-hour | APPBP2 | 1054.86873 | -0.1609514 | 0.03566953 | 0.00000649 | 0.04226332 |
| telencephalon | (left+right) | 24-hour vs 6-hour | KCNAB3 | 2585.9995 | 0.22187109 | 0.04876502 | 0.00000588 | 0.04226332 |

**Supplementary table 2E:** Differentially expressed genes (adjusted p-value of p<0.05) in the 6-hour light vs dark condition in the telencephalon (dark light used as baseline).

| **tissue** | **condition** | **comparison** | **gene** | **baseMean** | **log2FoldChange** | **lfcSE** | **pvalue** | **padj** |
|---|---|---|---|---|---|---|---|---|
| telencephalon | (left+right) | 6-hour vs dark | EIF3G | 1160.34631 | -0.1506437 | 0.03132693 | 7.51E-08 | 0.00051806 |
| telencephalon | (left+right) | 6-hour vs dark | PPIA | 13196.0459 | -0.126694 | 0.02661003 | 1.03E-07 | 0.00051806 |
| telencephalon | (left+right) | 6-hour vs dark | SLC25A11 | 1451.56744 | -0.1572396 | 0.035721 | 4.22E-07 | 0.00141733 |
| telencephalon | (left+right) | 6-hour vs dark | ATP5F1B | 12775.8482 | -0.0801506 | 0.01914887 | 0.00000255 | 0.00587467 |
| telencephalon | (left+right) | 6-hour vs dark | TIMP2 | 1604.08926 | -0.1290696 | 0.03216491 | 0.00000292 | 0.00587467 |
| telencephalon | (left+right) | 6-hour vs dark | HSPBP1 | 681.548488 | -0.154864 | 0.04181873 | 0.00000962 | 0.01411972 |
| telencephalon | (left+right) | 6-hour vs dark | UBE2S | 733.729652 | -0.1425134 | 0.03848105 | 0.00000981 | 0.01411972 |
| telencephalon | (left+right) | 6-hour vs dark | CSDC2 | 2118.32427 | -0.1302977 | 0.03584985 | 0.0000162 | 0.01815635 |
| telencephalon | (left+right) | 6-hour vs dark | LOC100857843 | 364.788507 | -0.1819671 | 0.05083283 | 0.0000153 | 0.01815635 |
| telencephalon | (left+right) | 6-hour vs dark | PSMD3 | 1932.32549 | -0.100968 | 0.02839807 | 0.0000205 | 0.02068155 |
| telencephalon | (left+right) | 6-hour vs dark | ARL5A | 2246.20331 | 0.11025665 | 0.03123875 | 0.0000236 | 0.02157185 |

| | | | | | | | | |
|---|---|---|---|---|---|---|---|---|
| telencephalon | (left+right) | 6-hour vs dark | MYL6 | 2703.09622 | -0.1219079 | 0.03646677 | 0.0000357 | 0.02993774 |
| telencephalon | (left+right) | 6-hour vs dark | ACTC2L | 20706.8203 | -0.1696576 | 0.05216149 | 0.0000412 | 0.03195625 |
| telencephalon | (left+right) | 6-hour vs dark | SLC30A10 | 1008.62434 | 0.12757973 | 0.03865467 | 0.0000483 | 0.03473452 |
| telencephalon | (left+right) | 6-hour vs dark | LRRC73 | 2609.3385 | -0.1313898 | 0.04057472 | 0.0000564 | 0.03553227 |
| telencephalon | (left+right) | 6-hour vs dark | PTPN11 | 3108.01458 | 0.13153429 | 0.04061689 | 0.0000545 | 0.03553227 |
| telencephalon | (left+right) | 6-hour vs dark | GDF3 | 2131.56141 | -0.1169233 | 0.03718755 | 0.0000733 | 0.04343434 |
| telencephalon | (left+right) | 6-hour vs dark | ATP11B | 2240.66765 | 0.12632601 | 0.04086115 | 0.0000904 | 0.04617023 |
| telencephalon | (left+right) | 6-hour vs dark | LOC112531384 | 304.781066 | -0.1753251 | 0.05763497 | 0.0000917 | 0.04617023 |
| telencephalon | (left+right) | 6-hour vs dark | SLC4A4 | 3626.77561 | 0.11603006 | 0.03726756 | 0.0000873 | 0.04617023 |
| telencephalon | (left+right) | 6-hour vs dark | HH3L | 1038.39347 | -0.1273351 | 0.04189216 | 0.0001057 | 0.04930644 |
| telencephalon | (left+right) | 6-hour vs dark | RAB40C | 378.300863 | -0.1477896 | 0.04934788 | 0.00011259 | 0.04930644 |
| telencephalon | (left+right) | 6-hour vs dark | VPS4A | 1268.45727 | -0.0916336 | 0.02967708 | 0.00010891 | 0.04930644 |

**Supplementary Table 3.** The table displays the enriched terms in the gene enrichment analysis indicating: Gene ontology (GO) for molecular function (MF), biological processes (BP) and cellular component (CC), or Kyoto Encyclopedia of Genes and Genomes (KEGG); term name; term ID; adjusted p value; negative log10 of adjusted value; term size; query size; intersection size; effective domain size and intersecting genes.

| source | Term name | Term id | Adjusted p value | Negative log10 of adjusted value | Term size | Query size | Inter-section size | Effective domain size | Intersections |
|---|---|---|---|---|---|---|---|---|---|
| GO:MF | oxygen carrier activity | GO:0005344 | 7.4895E-06 | 5.12554444 | 3 | 52 | 3 | 13050 | HBAD,HBBA,HBA1 |
| GO:MF | haptoglobin binding | GO:0031720 | 7.4895E-06 | 5.12554444 | 3 | 52 | 3 | 13050 | HBAD,HBBA,HBA1 |
| GO:MF | metal ion binding | GO:0046872 | 1.601E-05 | 4.79560693 | 1419 | 52 | 20 | 13050 | HBAD,ANGPT2,ADGRL4,FBLN2,HBBA,HBA1,CDH5,EHD2,EGFL7,MMP2,C9ORF58,CPM,ENPP2,ANPEP,CPA6,CPXM1,CA4,PRCP,CD93,TES |
| GO:MF | cation binding | GO:0043169 | 2.3151E-05 | 4.63543888 | 1477 | 52 | 20 | 13050 | HBAD,ANGPT2,ADGRL4,FBLN2,HBBA,HBA1,CDH5,EHD2,EGFL7,MMP2,C9ORF58,CPM,ENPP2,ANPEP,CPA6,CPXM1,CA4,PRCP,CD93,TES |
| GO:MF | oxygen binding | GO:0019825 | 0.00024743 | 3.60655294 | 9 | 52 | 3 | 13050 | HBAD,HBBA,HBA1 |
| GO:MF | platelet-derived growth factor binding | GO:0048407 | 0.00029373 | 3.53205631 | 10 | 52 | 3 | 13050 | COL1A1,PDGFRB,COL4A1 |
| GO:MF | ion binding | GO:0043167 | 0.00092905 | 3.03196304 | 2702 | 52 | 24 | 13050 | HBAD,ANGPT2,PDGFRB,ADGRL4,FBLN2,HBBA,HBA1,CDH5,EHD2,EGFL7,MMP2,TIE1,CHST15,ABCB1,C9ORF58,CPM,ENPP2,ANPEP,CPA6,CPXM1,CA4,PRCP,CD93,TES |
| GO:MF | metallocarboxypeptidase activity | GO:0004181 | 0.00092905 | 3.03196304 | 16 | 52 | 3 | 13050 | CPM,CPA6,CPXM1 |
| GO:MF | transition metal ion binding | GO:0046914 | 0.00092905 | 3.03196304 | 628 | 52 | 11 | 13050 | HBAD,HBA1,MMP2,CPM,ENPP2,ANPEP,CPA6,CPXM1,CA4,PRCP,TES |
| GO:MF | exopeptidase activity | GO:0008238 | 0.00172154 | 2.76408184 | 56 | 52 | 4 | 13050 | CPM,CPA6,CPXM1,PRCP |
| GO:MF | carboxypeptidase activity | GO:0004180 | 0.00172154 | 2.76408184 | 21 | 52 | 3 | 13050 | CPM,CPA6,CPXM1 |
| GO:MF | metallopeptidase activity | GO:0008237 | 0.00292919 | 2.53325272 | 126 | 52 | 5 | 13050 | MMP2,CPM,ANPEP,CPA6,CPXM1 |
| GO:MF | peroxidase activity | GO:0004601 | 0.00339128 | 2.4696365 | 28 | 52 | 3 | 13050 | HBAD,HBBA,HBA1 |
| GO:MF | calcium ion binding | GO:0005509 | 0.00339128 | 2.4696365 | 411 | 52 | 8 | 13050 | ADGRL4,FBLN2,CDH5,EHD2,EGFL7,C9ORF58,ENPP2,CD93 |
| GO:MF | extracellular matrix structural constituent | GO:0005201 | 0.00339128 | 2.4696365 | 29 | 52 | 3 | 13050 | COL1A1,COL4A1,DCN |
| GO:MF | oxidoreductase activity, acting on peroxide as acceptor | GO:0016684 | 0.00388915 | 2.41014571 | 31 | 52 | 3 | 13050 | HBAD,HBBA,HBA1 |
| GO:MF | collagen binding | GO:0005518 | 0.00471541 | 2.32648072 | 35 | 52 | 3 | 13050 | VWF,DCN,ITGA2 |
| GO:MF | metalloexopeptidase activity | GO:0008235 | 0.00471541 | 2.32648072 | 35 | 52 | 3 | 13050 | CPM,CPA6,CPXM1 |
| GO:MF | molecular carrier activity | GO:0140104 | 0.00471541 | 2.32648072 | 34 | 52 | 3 | 13050 | HBAD,HBBA,HBA1 |
| GO:MF | zinc ion binding | GO:0008270 | 0.00624548 | 2.20443386 | 470 | 52 | 8 | 13050 | MMP2,CPM,ENPP2,ANPEP,CPA6,CPXM1,CA4,TES |
| GO:MF | binding | GO:0005488 | 0.0067878 | 2.16827104 | 7635 | 52 | 42 | 13050 | HBAD,APCDD1,ANGPT2,COL1A1,PDGFRB,ADGRL4,FBLN2,KANK3,HBBA,HBA1,SHE,CDH5,SDCBP,VWF,ACTA2,COL4A1,EHD2,GDF10,EGFL7,MMP2,TIE1,FLI1,MSN,TRIL, |

| | | | | | | | | | |
|---|---|---|---|---|---|---|---|---|---|
| | | | | | | | | | CHST15,DIO2,VIM,ABCB1,DCN,C9ORF58,CPM,ENPP2,ANPEP,CPA6,CPXM1,ITGA2,CA4,TMPRSS9,PAX2,PRCP,CD93,TES |
| GO:MF | peptidase activity | GO:0008233 | 0.00742163 | 2.12950095 | 373 | 52 | 7 | 13050 | MMP2,CPM,ANPEP,CPA6,CPXM1,TMPRSS9,PRCP |
| GO:MF | antioxidant activity | GO:0016209 | 0.00821156 | 2.08557418 | 45 | 52 | 3 | 13050 | HBAD,HBBA,HBA1 |
| GO:BP | oxygen transport | GO:0015671 | 0.00075453 | 3.12232489 | 6 | 52 | 3 | 13050 | HBAD,HBBA,HBA1 |
| GO:BP | gas transport | GO:0015669 | 0.00075453 | 3.12232489 | 7 | 52 | 3 | 13050 | HBAD,HBBA,HBA1 |
| GO:BP | vasculature development | GO:0001944 | 0.00075453 | 3.12232489 | 387 | 52 | 10 | 13050 | ANGPT2,COL1A1,CDH5,ACTA2,COL4A1,MMP2,TIE1,DCN,ENPP2,PRCP |
| GO:BP | blood vessel development | GO:0001568 | 0.00075453 | 3.12232489 | 364 | 52 | 10 | 13050 | ANGPT2,COL1A1,CDH5,ACTA2,COL4A1,MMP2,TIE1,DCN,ENPP2,PRCP |
| GO:BP | hydrogen peroxide catabolic process | GO:0042744 | 0.00081654 | 3.08802238 | 8 | 52 | 3 | 13050 | HBAD,HBBA,HBA1 |
| GO:BP | anatomical structure formation involved in morphogenesis | GO:0048646 | 0.00199355 | 2.70037274 | 567 | 52 | 11 | 13050 | ANGPT2,COL1A1,CDH5,COL4A1,MMP2,TIE1,DCN,ENPP2,ITGA2,PAX2,PRCP |
| GO:BP | regulation of cell migration | GO:0030334 | 0.00199355 | 2.70037274 | 460 | 52 | 10 | 13050 | ANGPT2,COL1A1,PDGFRB,CDH5,SDCBP,TIE1,MSN,DCN,ENPP2,PRCP |
| GO:BP | cell migration | GO:0016477 | 0.00246356 | 2.60843728 | 714 | 52 | 12 | 13050 | ANGPT2,COL1A1,PDGFRB,CDH5,SDCBP,TIE1,MSN,DCN,ENPP2,ITGA2,PAX2,PRCP |
| GO:BP | regulation of cell motility | GO:2000145 | 0.00246356 | 2.60843728 | 485 | 52 | 10 | 13050 | ANGPT2,COL1A1,PDGFRB,CDH5,SDCBP,TIE1,MSN,DCN,ENPP2,PRCP |
| GO:BP | hydrogen peroxide metabolic process | GO:0042743 | 0.00260926 | 2.5834821 | 14 | 52 | 3 | 13050 | HBAD,HBBA,HBA1 |
| GO:BP | regulation of cellular component movement | GO:0051270 | 0.00297639 | 2.52631067 | 513 | 52 | 10 | 13050 | ANGPT2,COL1A1,PDGFRB,CDH5,SDCBP,TIE1,MSN,DCN,ENPP2,PRCP |
| GO:BP | regulation of locomotion | GO:0040012 | 0.00297639 | 2.52631067 | 508 | 52 | 10 | 13050 | ANGPT2,COL1A1,PDGFRB,CDH5,SDCBP,TIE1,MSN,DCN,ENPP2,PRCP |
| GO:BP | cell motility | GO:0048870 | 0.00404488 | 2.39309457 | 787 | 52 | 12 | 13050 | ANGPT2,COL1A1,PDGFRB,CDH5,SDCBP,TIE1,MSN,DCN,ENPP2,ITGA2,PAX2,PRCP |
| GO:BP | localization of cell | GO:0051674 | 0.00404488 | 2.39309457 | 787 | 52 | 12 | 13050 | ANGPT2,COL1A1,PDGFRB,CDH5,SDCBP,TIE1,MSN,DCN,ENPP2,ITGA2,PAX2,PRCP |
| GO:BP | circulatory system development | GO:0072359 | 0.00404488 | 2.39309457 | 546 | 52 | 10 | 13050 | ANGPT2,COL1A1,CDH5,ACTA2,COL4A1,MMP2,TIE1,DCN,ENPP2,PRCP |
| GO:BP | angiogenesis | GO:0001525 | 0.004662 | 2.33142741 | 253 | 52 | 7 | 13050 | ANGPT2,CDH5,COL4A1,TIE1,DCN,ENPP2,PRCP |
| GO:BP | cell-substrate adhesion | GO:0031589 | 0.00555251 | 2.25551048 | 180 | 52 | 6 | 13050 | ANGPT2,COL1A1,FBLN2,VWF,MMP2,ITGA2 |
| GO:BP | regulation of vasculature development | GO:1901342 | 0.00736465 | 2.13284808 | 126 | 52 | 5 | 13050 | ANGPT2,CDH5,TIE1,DCN,ENPP2 |
| GO:BP | reactive oxygen species metabolic process | GO:0072593 | 0.00736465 | 2.13284808 | 121 | 52 | 5 | 13050 | HBAD,HBBA,HBA1,PAX2,PRCP |
| GO:BP | cellular oxidant detoxification | GO:0098869 | 0.00736465 | 2.13284808 | 24 | 52 | 3 | 13050 | HBAD,HBBA,HBA1 |
| GO:BP | bone trabecula formation | GO:0060346 | 0.00736465 | 2.13284808 | 5 | 52 | 2 | 13050 | COL1A1,MMP2 |
| GO:BP | direct ossification | GO:0036072 | 0.00736465 | 2.13284808 | 5 | 52 | 2 | 13050 | COL1A1,MMP2 |

| | | | | | | | | | |
|---|---|---|---|---|---|---|---|---|---|
| GO:BP | blood vessel maturation | GO:0001955 | 0.00736465 | 2.13284808 | 5 | 52 | 2 | 13050 | CDH5,MMP2 |
| GO:BP | regulation of angiogenesis | GO:0045765 | 0.00736465 | 2.13284808 | 124 | 52 | 5 | 13050 | ANGPT2,CDH5,TIE1,DCN,ENPP2 |
| GO:BP | locomotion | GO:0040011 | 0.00736465 | 2.13284808 | 884 | 52 | 12 | 13050 | ANGPT2,COL1A1,PDGFRB,CDH5,SDCBP,TIE1,MSN,DCN,ENPP2,ITGA2,PAX2,PRCP |
| GO:BP | intramembranous ossification | GO:0001957 | 0.00736465 | 2.13284808 | 5 | 52 | 2 | 13050 | COL1A1,MMP2 |
| GO:BP | external encapsulating structure organization | GO:0045229 | 0.00855748 | 2.06765387 | 136 | 52 | 5 | 13050 | COL1A1,FBLN2,COL4A1,MMP2,TIE1 |
| GO:BP | extracellular structure organization | GO:0043062 | 0.00855748 | 2.06765387 | 136 | 52 | 5 | 13050 | COL1A1,FBLN2,COL4A1,MMP2,TIE1 |
| GO:BP | extracellular matrix organization | GO:0030198 | 0.00855748 | 2.06765387 | 136 | 52 | 5 | 13050 | COL1A1,FBLN2,COL4A1,MMP2,TIE1 |
| GO:BP | blood vessel morphogenesis | GO:0048514 | 0.00880723 | 2.05516061 | 310 | 52 | 7 | 13050 | ANGPT2,CDH5,COL4A1,TIE1,DCN,ENPP2,PRCP |
| GO:BP | cellular detoxification | GO:1990748 | 0.00897485 | 2.04697276 | 30 | 52 | 3 | 13050 | HBAD,HBBA,HBA1 |
| GO:BP | collagen metabolic process | GO:0032963 | 0.00959891 | 2.01777813 | 31 | 52 | 3 | 13050 | COL1A1,MMP2,VIM |
| GO:CC | haptoglobin-hemoglobin complex | GO:0031838 | 1.0384E-05 | 4.98363892 | 3 | 52 | 3 | 13050 | HBAD,HBBA,HBA1 |
| GO:CC | cell periphery | GO:0071944 | 1.3806E-05 | 4.8599252 | 2226 | 52 | 25 | 13050 | APCDD1,SLC28A3,ANGPT2,COL1A1,PDGFRB,SSPN,ADGRL4,FBLN2,CDH5,SDCBP,VWF,COL4A1,GYPC,MMP2,TIE1,GPER1,MSN,VIM,ABCB1,C9ORF58,ANPEP,ITGA2,CA4,CD93,TES |
| GO:CC | hemoglobin complex | GO:0005833 | 1.3806E-05 | 4.8599252 | 4 | 52 | 3 | 13050 | HBAD,HBBA,HBA1 |
| GO:CC | collagen-containing extracellular matrix | GO:0062023 | 0.00012285 | 3.91061731 | 101 | 52 | 6 | 13050 | ANGPT2,COL1A1,FBLN2,VWF,COL4A1,MMP2 |
| GO:CC | extracellular space | GO:0005615 | 0.00077093 | 3.11298721 | 394 | 52 | 9 | 13050 | ANGPT2,COL1A1,HBBA,HBA1,GDF10,MMP2,DCN,ENPP2,CA4 |
| GO:CC | extracellular region | GO:0005576 | 0.00077483 | 3.1107909 | 622 | 52 | 11 | 13050 | ANGPT2,COL1A1,FBLN2,HBBA,HBA1,VWF,GDF10,MMP2,DCN,ENPP2,CA4 |
| GO:CC | extracellular matrix | GO:0031012 | 0.00090251 | 3.04455012 | 162 | 52 | 6 | 13050 | ANGPT2,COL1A1,FBLN2,VWF,COL4A1,MMP2 |
| GO:CC | external encapsulating structure | GO:0030312 | 0.00090251 | 3.04455012 | 162 | 52 | 6 | 13050 | ANGPT2,COL1A1,FBLN2,VWF,COL4A1,MMP2 |
| GO:CC | collagen trimer | GO:0005581 | 0.00090251 | 3.04455012 | 18 | 52 | 3 | 13050 | COL1A1,COL4A1,COL9A3 |
| GO:CC | plasma membrane | GO:0005886 | 0.00095116 | 3.02174586 | 2039 | 52 | 20 | 13050 | APCDD1,SLC28A3,PDGFRB,SSPN,ADGRL4,CDH5,SDCBP,GYPC,MMP2,TIE1,GPER1,MSN,VIM,ABCB1,C9ORF58,ANPEP,ITGA2,CA4,CD93,TES |
| GO:CC | cell surface | GO:0009986 | 0.00175915 | 2.75469654 | 279 | 52 | 7 | 13050 | CDH5,MSN,ABCB1,ANPEP,ITGA2,CA4,CD93 |
| GO:CC | complex of collagen trimers | GO:0098644 | 0.00468189 | 2.32957849 | 7 | 52 | 2 | 13050 | COL1A1,COL4A1 |
| GO:CC | external side of plasma membrane | GO:0009897 | 0.00570493 | 2.24374983 | 89 | 52 | 4 | 13050 | CDH5,ANPEP,ITGA2,CA4 |
| GO:CC | focal adhesion | GO:0005925 | 0.00989633 | 2.00452602 | 105 | 52 | 4 | 13050 | MSN,C9ORF58,ITGA2,TES |
| KEGG | ECM-receptor interaction | KEGG:04512 | 0.00029938 | 3.5237798 | 71 | 52 | 5 | 13050 | COL1A1,VWF,COL4A1,COL9A3,ITGA2 |
| KEGG | Focal adhesion | KEGG:04510 | 0.00088241 | 3.05432838 | 168 | 52 | 6 | 13050 | COL1A1,PDGFRB,VWF,COL4A1,COL9A3,ITGA2 |

**Supplementary Table 4.** Number of pair reads per sample

| Sample | Pair reads per sample |
|---|---|
| WTCHG_328669_209 | 6921213 |
| WTCHG_328669_210 | 4019341 |
| WTCHG_328669_221 | 4004936 |
| WTCHG_328669_222 | 12728700 |
| WTCHG_328669_223 | 14060200 |
| WTCHG_328669_224 | 15842389 |
| WTCHG_328669_241 | 17379215 |
| WTCHG_328669_242 | 19239117 |
| WTCHG_328669_243 | 11350328 |
| WTCHG_328669_244 | 10017003 |
| WTCHG_328669_245 | 11485751 |
| WTCHG_328669_246 | 4245942 |
| WTCHG_328669_247 | 6415919 |
| WTCHG_328669_248 | 13312943 |
| WTCHG_328669_249 | 7824585 |
| WTCHG_328669_250 | 14705126 |
| WTCHG_328669_273 | 18767526 |
| WTCHG_328669_274 | 3353362 |
| WTCHG_328669_275 | 11265358 |
| WTCHG_328669_276 | 14875611 |
| WTCHG_328669_277 | 3338442 |
| WTCHG_328669_278 | 12290015 |
| WTCHG_328669_279 | 3753295 |
| WTCHG_328669_280 | 16741498 |
| WTCHG_328669_289 | 6731848 |
| WTCHG_328669_290 | 14389416 |
| WTCHG_328669_291 | 4171235 |
| WTCHG_328669_292 | 15825416 |
| WTCHG_328669_293 | 12529431 |
| WTCHG_328669_294 | 11315392 |
| WTCHG_328670_211 | 4353541 |
| WTCHG_328670_212 | 4325944 |
| WTCHG_328670_213 | 10318631 |
| WTCHG_328670_214 | 4950807 |
| WTCHG_328670_215 | 13237564 |
| WTCHG_328670_216 | 16354197 |
| WTCHG_328670_217 | 19189283 |
| WTCHG_328670_218 | 14401929 |
| WTCHG_328670_219 | 4473086 |
| WTCHG_328670_220 | 5078432 |
| WTCHG_328670_233 | 7834320 |
| WTCHG_328670_234 | 18453221 |

| | |
|---|---|
| WTCHG_328670_235 | 17376052 |
| WTCHG_328670_236 | 9388101 |
| WTCHG_328670_237 | 13398756 |
| WTCHG_328670_238 | 13515330 |
| WTCHG_328670_239 | 6772601 |
| WTCHG_328670_240 | 6723865 |
| WTCHG_328670_251 | 4590925 |
| WTCHG_328670_252 | 5049950 |
| WTCHG_328670_253 | 21780719 |
| WTCHG_328670_254 | 16920263 |
| WTCHG_328670_255 | 4791542 |
| WTCHG_328670_256 | 5950143 |
| WTCHG_328670_257 | 22089522 |
| WTCHG_328670_258 | 4620560 |
| WTCHG_328670_259 | 9967688 |
| WTCHG_328670_295 | 19304576 |
| WTCHG_328670_296 | 10406265 |
| WTCHG_328670_301 | 11198668 |
